# TRPV1 Defends the Healthy Murine Cornea against *Staphylococcus aureus* Adhesion Independently of Sensory Nerve Firing

**DOI:** 10.1101/2025.04.12.648450

**Authors:** Orneika Flandrin, Yujia Yang, Sara Abboud, Naren G Kumar, Ananya Datta, Eric Jedel, Diana Bautista, David Evans, Suzanne Fleiszig

**Author notes:** **Correspondence.** Dr. Suzanne M. J. Fleiszig, Herbert Wertheim School of Optometry & Vision Science, University of California, Berkeley CA 94720, USA. **Present address.** New England College of Optometry. Boston, MA 02115, USA. **Commercial relationships**. None to declare for each author.

## Abstract

**Purpose:** Previously we showed that transient receptor potential ion channels TRPA1 and TRPV1 selectively protect the cornea against bacterial adhesion, with TRPA1 countering the Gram-negative pathogen *Pseudomonas aeruginosa* and TRPV1 countering environmental bacteria. Here, we explored parameters of this specificity using a Gram-positive pathogen *Staphylococcus aureus*.

**Methods:** Healthy corneas of C57BL/6J wild-type (WT), TRPA1 (-/-) or TRPV1 (-/-) mice were challenged with *S. aureus* for 4 or 6 hours. Some experiments instead/also used resiniferatoxin (RTX) to deplete TRPV1-expressing nerves, JNJ-17203212 to selectively antagonize TRPV1, or the anesthetic bupivacaine to inhibit nerve firing. Adherent bacteria were quantified using FISH labeling (16S rRNA-targeted probe). Lyz2^+^, CD11c^+^ and CD45^+^ cells were visualized/quantified using hybrid mT/mG + LysMcre mice (red cell membranes; Lyz2^+^-GFP), CD11c^+^-YFP mice, and anti-CD45-antibody respectively.

**Results:** Corneas of TRPV1 (-/-) not TRPA1 (-/-) mice were found more susceptible to *S. aureus* adhesion compared to WT. Accordingly, either ablation of TRPV1-expressing nerves or TRPV1 antagonism increased adhesion. Defense against *S. aureus* adhesion did not depend on nerve firing. Despite having no significant impact on CD11c^+^ or Lyz2^+^ cell numbers, *S. aureus* challenge increased CD45^+^ cell counts, also dependent on TRPV1-expressing nerves, and it increased Lyz2^+^ cell sphericity and volume.

**Conclusion:** Healthy corneas utilize TRPV1 to protect against *S. aureus* adhesion independently of sensory nerve firing. This contrasts with defense against *P. aeruginosa* adhesion which requires TRPA1 and nerve firing. How the differential immune cell responses to these two pathogens relate to TRP-dependent defense against adhesion remains to be determined.

## Introduction

Despite regular exposure to environmental microbes, healthy corneas lack a microbiome and exhibit a unique resistance to colonization by prominent pathogens, including *Pseudomonas aeruginosa* (Gram-negative) and *Staphylococcus aureus* (Gram-positive), the latter existing as a skin commensal and also a common cause of ocular infections ^1–6^. Understanding the healthy cornea’s ability to avoid colonization, an important step in infection pathogenesis, is essential to preventing sight-threatening infections.

Previous work by our laboratory and others has highlighted an important role for numerous factors in mediating the exceptional intrinsic defense of the healthy cornea against *P. aeruginosa, S. aureus* and other bacteria. These factors include; mucus ^7^, secretory IgA ^8^, surfactant protein-D^9^, the basement membrane ^10^, surface mucins ^11,12^, tight junctions ^13^, antimicrobial peptides ^3,14–17^ including cytokeratins^18,19^, DMBT1^20^, resident immune cells ^21^ and signaling molecules including MyD88 and IL-1R^1,22–24^.

More recently, we discovered that sensory nerves, abundant in the cornea, play a role in defending against *P. aeruginosa* adhesion ^25^. Sensory nerves, along with their polymodal transient receptor potential (TRP) ion channels including TRPA1 (Ankyrin) and TRPV1 (Vanilloid), are known to detect harmful factors including bacterial ligands or toxins with an induction of pain, inflammation and immune response activation and modulation ^26–34^. Our study showed that TRPA1 prevented *P. aeruginosa* adhesion to a healthy cornea correlating with CD45+ immune cell recruitment ^25^. Additionally, corneal sensory nerve firing, inhibited by bupivacaine anesthesia, was also necessary for defending healthy and superficially-injured corneas against *P. aeruginosa*, correlating with a sensory nerve-dependent CD11c^+^ cell immune response ^25^, that we previously showed could inhibit *P. aeruginosa* adhesion to the cornea after superficial-injury ^21^. However, TRPV1-deficient corneas showed increased colonization by environmental bacteria, most likely Gram-positive in nature ^25^.

Here, we sought to determine if TRPA1 and TRPV1 exert pathogen specificity in defending the healthy cornea against bacterial adhesion by using a different ocular pathogen *Staphylococcus aureus*. Bacterial adhesion and immune cell responses to *S. aureus* challenge were examined using TRPA1 (-/-) and TRPV1 (-/-) mice combined with pharmacological inhibition of receptors and sensory nerve firing. Comparisons with *P. aeruginosa* were included as applicable.

## Materials and Methods

### Mice

Six-to twelve-week-old male or female C57BL/6J wild-type (WT) mice (Jackson Laboratory) and gene knockouts in TRPV1 (-/-) or TRPA1 (-/-) (kindly provided by Dr. Diana Bautista, University of California, Berkeley) were used for bacterial adhesion experiments. In some experiments, transgenic mT/mG + LysMcre hybrid WT mice, i.e. a mT/mG mouse (red cell membranes, Jackson Laboratory) crossed with a LysMcre mouse (green Lyz2^+^ cells, Jackson Laboratory), or CD11c^+^- YFP (yellow CD11c^+^ cells, Jackson Laboratory) mice were used to assess immune cell responses to bacteria. Videos to demonstrate Imaris immune cell processing methods used a mT/mG + LysMcre mouse. All procedures were carried out per standards established by the Association for the Research in Vision and Ophthalmology (ARVO), under protocol AUP-2019-06-12322 approved by the Animal Care and Use Committee, the University of California Berkeley (an AAALAC-accredited institution). The protocol adheres to PHS policy on the humane care and use of laboratory animals, and the guide for the care and use of laboratory animals.

### Bacterial Strains

*Staphylococcus aureus* S33, a clinical isolate from a human ocular infection, was used throughout the study. In some experiments, *Pseudomonas aeruginosa* PAO1 (tdTomato) was also used for comparison to our prior work ^25^. Bacteria were prepared by growth on a trypticase soy agar plate overnight for ∼16 hours at 37°C, followed by suspension in phosphate-buffered saline (PBS) to a concentration of ∼10^11^ colony-forming units (CFU)/ml.

### Bacterial Adhesion Assay using Fluorescence *in Situ* Hybridization (FISH)

An *in vivo* model of bacterial adhesion was used as we have described previously ^2,25^. Mice were anesthetized by intraperitoneal injection of ketamine (80 - 100 mg/Kg) and dexmedetomidine (0.25-0.5 mg/Kg). Corneas were inoculated with 5 μl of bacterial suspension once every hour for 4 hours (4 inoculations) while contralateral corneas were sham-inoculated with PBS. Animals remained anesthetized on a heated pad for the duration of bacterial exposure. After 4 hours, animals were euthanized by intraperitoneal injection of ketamine (80–100 mg/Kg) and xylazine (5-10 mg/Kg), or isoflurane (5 %) for 10 minutes, either followed by cervical dislocation. Eyes were enucleated, rinsed with PBS, and fixed in 2 % paraformaldehyde overnight at 4°C. For *ex vivo* experiments, freshly enucleated eyes were submerged in 200 μl of bacterial suspension for 6 hours at 37°C followed by a PBS rinse with fixation.

Fixed whole eyes were labeled for adherent bacteria using a universal 16S rRNA-targeted FISH probe as previously described ^1,25,35^. Briefly, fixed eyes were washed in PBS, 80 % ethanol, and 95 % ethanol for 10 minutes each at room temperature. Eyes were placed in hybridization buffer (0.9 M NaCl, 20 mM Tris-HCl, and 0.01 % SDS) followed by incubation at 55°C for 30 minutes. The 16S rRNA-targeted gene probe [Alexa488]-GCTGCCTCCCGTAGGAGT- [Alexa488] (Eurofins Genomics) was added to eyes to a final concentration of 100 nM for overnight incubation at 55°C. Adherent bacteria were imaged and quantified as described below.

### TRPA1/TRPV1 Nociceptor Ablation and Corneal Sensory Nerve Block

TRPV1-expressing corneal nerves were selectively ablated using resiniferatoxin (RTX) (AdipoGen: AG-CN2-0534-MC05) as previously described ^25,36,37^. Briefly, mice were lightly anesthetized with isoflurane (3 %), and 100 μl RTX solution was injected subcutaneously in the scruff of the neck at a final concentration of 30 μM for 3 consecutive days. To inhibit corneal sensory nerve firing, 0.5 % bupivacaine hydrochloride solution was injected into the subconjunctival sac (5 μl) and added topically (5 μl) in anesthetized mice for 20 minutes ^25^. Following RTX ablation or sensory nerve block, bacterial adherence was assessed as above.

### TRPV1 Ion Channel Inhibition

The JNJ-17203212 (Cayman Chemical; #30930) antagonist was used to selectively inhibit TRPV1 channel activity as previously described ^38,39^. Briefly, mice were anesthetized and 5 μl antagonist was injected into the subconjunctival sac at a final concentration of 500 μM. An additional 5 μl was immediately added topically. After 20 minutes, corneas were washed with PBS followed by a bacterial adherence assay. In a control experiment without inoculation, a capsaicin eye wipe test was performed after antagonist treatment and determined the effective duration of TRPV1 block to be 3 hours. Corneal epithelial integrity was also assessed 4 hours after antagonist treatment by adding 5 μl of fluorescein solution (0.02 %) to the ocular surface followed by slit-lamp imaging.

### Immunohistochemistry

In some experiments, after the bacterial adhesion assay, enucleated eyes were fixed overnight in 2 % paraformaldehyde then washed for 10 minutes with rotation as previously described ^25,37^. Corneal dissections were performed under a dissecting microscope and dissected corneas placed in a blocking solution (3 % bovine serum albumin with 0.3 % Triton X-100 in PBS) for 1 hour at room temperature. Corneas were then incubated in primary antibody (rat anti-mouse CD45+ [1:500; BD Pharmingen: #550539]) overnight at 4°C with rotation. Corneas were transferred to secondary antibody (anti-rat antibody [Life Technologies: #A21434]) diluted in DAPI (4,6- diamidino-2-phenylindole dihydrochloride; 12.5 μg/mL; Thermo Fisher: #D1306) for 2 hours at room temperature with rotation and covered with aluminum foil. Corneas were transferred to fresh PBS and washed 3 times for 10 minutes with rotation at room temperature and flat-mounted with Prolong Gold (Thermo Fisher: #P36970) before confocal imaging.

### Confocal Microscopy and Image Analysis

Samples were imaged using an Olympus FV1000 confocal microscope with a 20×/ 1.0 NA water-dipping objective. The 488 nm laser was used for detection of bacteria labeled with the FISH probe, fluorescein, or Lyz2^+^-GFP cells. The 515 nm laser was used for CD11c^+^-YFP cells, 559 nm used for red cell membranes, and the 635 nm laser used for corneas labeled with anti-CD45 antibody. The 635 nm laser was also used to visualize corneal surface reflectance (excitation and emission at the same wavelength). Z stacks were acquired at a 0.4 μm step size and an aspect ratio of 1024 μm x 1024 μm for bacteria detection. Adherent bacteria were identified and quantified using Imaris spot detection. For immune cells, a 1μm step size and 512 μm x 512 μm aspect ratio were used. Acquired Z stacks were reconstructed as 3-D images using Imaris Software. Corneal immune cell morphology was analyzed as previously described ^37,40^, and the processes summarized in Supplemental Fig. S1. Supplemental Videos 1 and 2 show examples of the processes illustrated in Fig. S1: Video 1 showing surface rendering and distance transformation of corneal immune cells, Video 2 showing color-coding of immune cells based on their volume, sphericity and distance from the endothelium.

## Statistical Analysis

Statistical analysis was performed using Prism (GraphPad Software, Inc.). Data were expressed as the mean ± standard deviation (SD). An unpaired Student’s t-Test was used for two group comparisons. Comparisons between three or more groups were performed using One-way or Two-way ANOVA tests with Tukey’s multiple comparisons as indicated in each figure. P values < 0.05 were considered significant. All experiments were repeated at least twice.

## Results

### TRPV1 is Required for Corneal Defense versus *S. aureus* Adhesion

Our previous findings demonstrated a role for TRPA1, but not TRPV1, in preventing adhesion of *P. aeruginosa* to the healthy and superficially-injured murine cornea^25^. The same study showed that TRPV1 helped prevent corneal colonization by environmental bacteria. Here, we first tested whether TRPA1 or TRPV1 defended the murine cornea against adhesion of *S. aureus*. Experiments using gene knockout mice showed that TRPV1 was required for defending the healthy cornea against deliberately-inoculated *S. aureus*, with TRPA1 (-/-) mice showing no difference from wild-type with both resisting bacterial adhesion (Fig. 1A). Consistent with our previous work, there was an increase in adhesion of background environmental bacteria on sham-inoculated TRPV1 (-/-) corneas versus WT (not shown). Therefore, quantification of bacterial adhesion after *S. aureus* inoculation was normalized to background (baseline) and showed a 5.6-fold increase in *S. aureus* adhesion associated with TRPV1 (-/-) corneas versus WT (Fig. 1B, P < 0.05, Two-way ANOVA). Adherent bacteria remained surface-attached and did not penetrate the epithelium (Fig. 1C). Thus, TRPV1 is needed to prevent *S. aureus* adhesion to the healthy murine cornea, contrasting with the requirement for TRPA1 to defend against the adhesion of *P. aeruginosa*.

**Figure 1.**
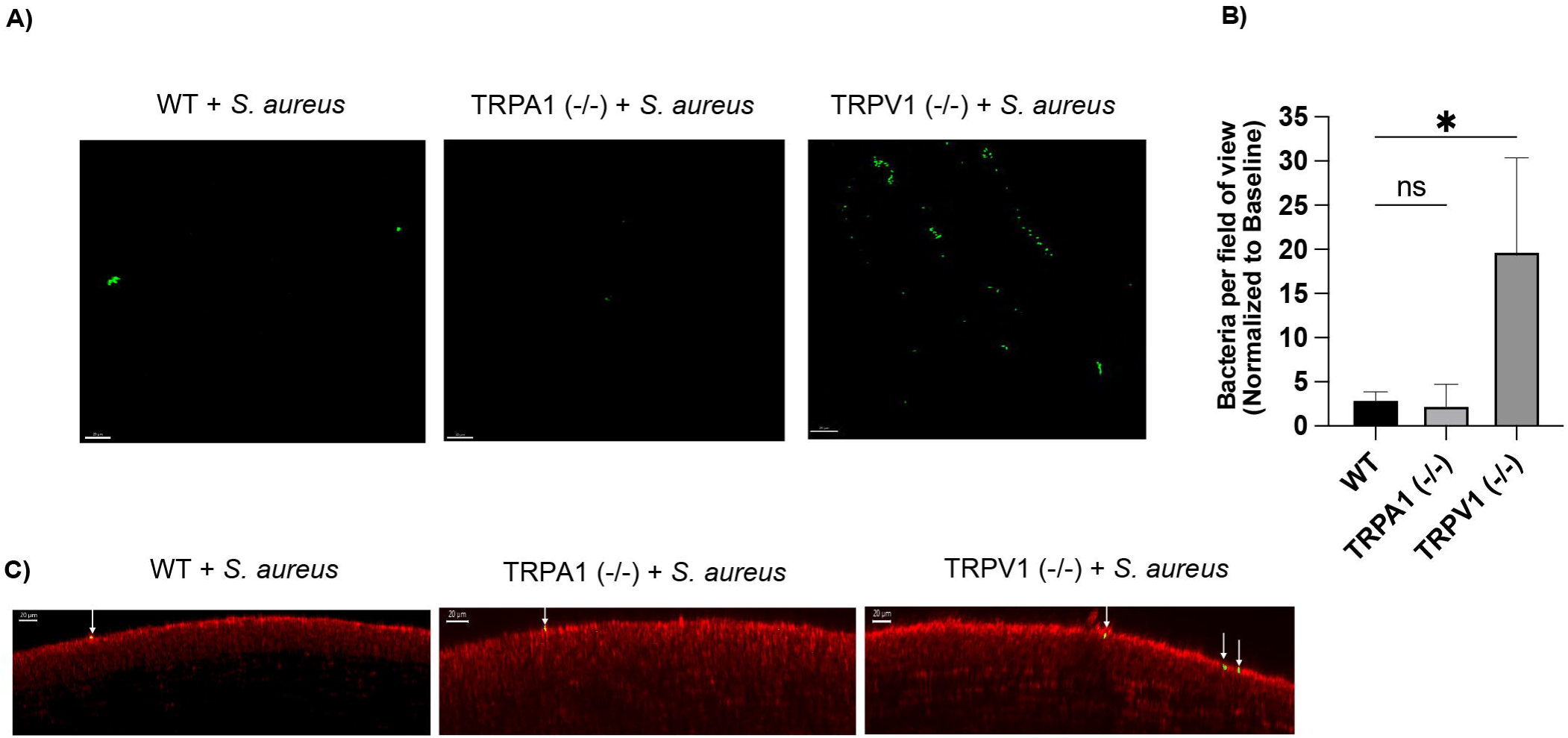
Corneal defense against *S. aureus* adhesion requires TRPV1 ion channels. **(A)** Corneas were labeled using a universal 16S rRNA-targeted FISH probe. Representative images showing bacteria (green) adherent to healthy wild-type (WT), TRPA1 (-/-) or TRPV1 (-/-) mouse corneas 4 hours after inoculation with ∼10^11^ CFU/ml *S. aureus*. 20X Objective. Scale bars = 20 μm. Only the bacterial channel shown. **(B)** Quantification shows significantly greater FISH-labeled bacteria adherent to TRPV1 (-/-) corneas at 4 hours after deliberate inoculation with *S. aureus* (5.6-fold) compared to those of WT or TRPA1 (-/-) mice. Data were normalized to baseline bacteria and expressed as the mean ± SD of bacteria per field of view. * P ˂ 0.05, ns = Not Significant (Two-way ANOVA with Tukey’s multiple comparisons). **(C)** XZ optical slices show that bacteria were only surface attached following *S. aureus* inoculation (green, indicated by white arrows) and did not penetrate the corneal epithelium (red). Scale bars = 20 μm.

### Corneal Defense vs. *S. aureus* Adhesion is Inhibited by Resiniferatoxin, Eliminated *Ex vivo* but Retained after Bupivacaine Treatment

To assess the role of TRPV1-associated sensory nerves in the corneal defense against *S. aureus* adhesion, we first used resiniferatoxin (RTX) to ablate TRPV1-expressing neurons, and those co-expressing TRPA1 ^41^. WT mice were treated with RTX for 3 days followed by inoculation with *S. aureus in vivo*. Fig. 2 shows effective RTX ablation of sensory nerves (Fig. 2A) associated with a significant increase in *S. aureus* adhesion, that was quantified as a 5.3-fold increase in *S. aureus* adhesion to the cornea versus wild-type after normalization to background environmental bacteria (baseline) (Fig. 2B, P < 0.05, Two-way ANOVA).

**Figure 2.**
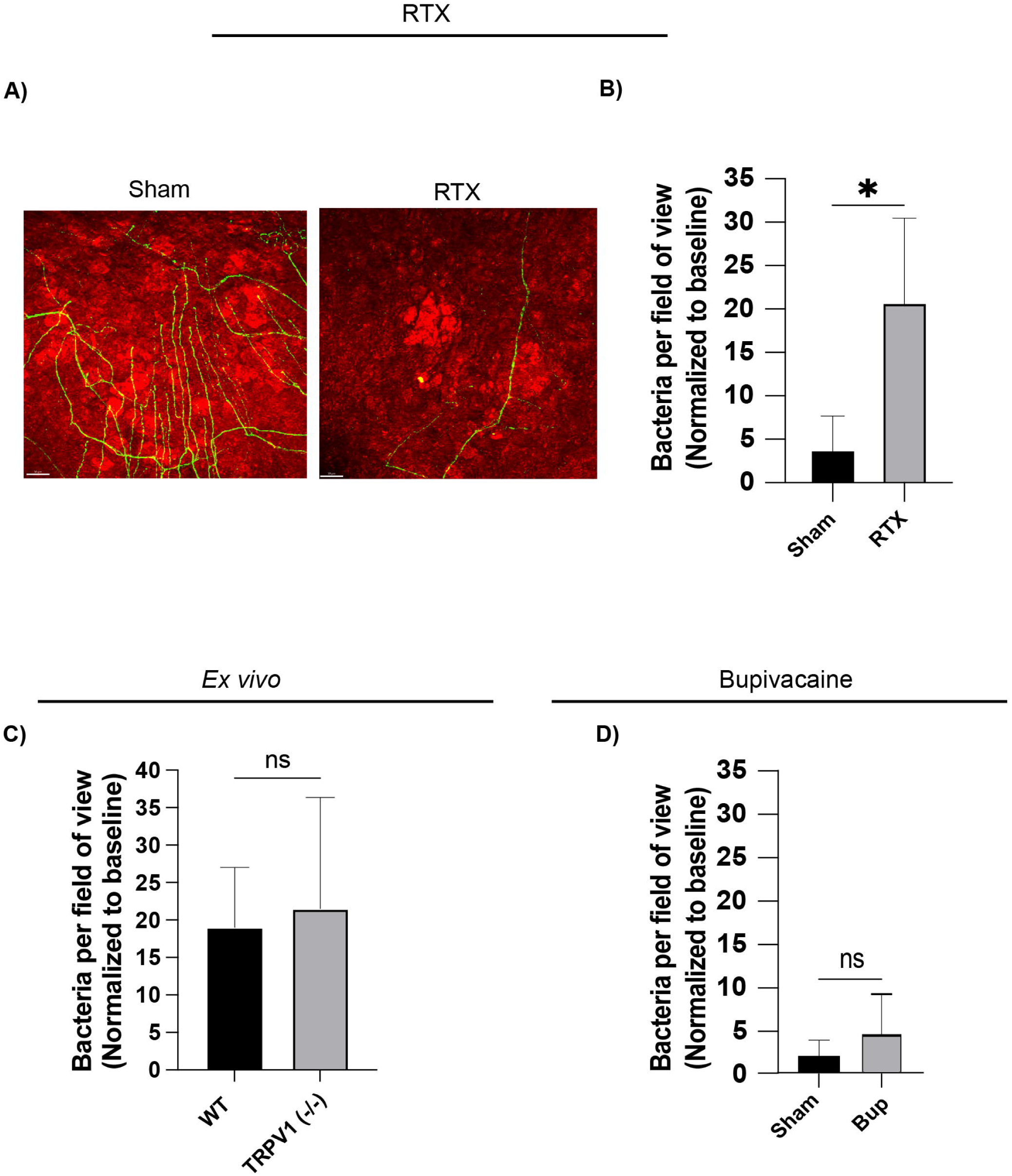
Corneal defense against *S. aureus* adhesion is inhibited by RTX treatment, absent *ex vivo*, but retained after bupivacaine treatment. **(A)** Corneal nerve density decreased following RTX (30 μM) ablation over 3 days, as shown by representative images (corneal nerves in green, corneal epithelium in red). Scale bars = 50 μm. **(B)** Quantification showing significantly greater FISH-labeled bacteria on RTX treated corneas at 4 hours after inoculation with *S. aureus* (5.3-fold) compared to WT mice. Data were normalized to baseline bacteria and expressed as the mean ± SD of bacteria per field of view. * P ˂ 0.05, ns = Not Significant (Two-way ANOVA with Tukey’s multiple comparisons). **(C)** *Ex vivo*, corneal defense against *S. aureus* adhesion was lost with wild-type corneas showing an increase in adhesion to match the TRPV1 (-/-) mice. Data were expressed as the mean ± SD of bacteria per field of view. ns = Not Significant (Two-way ANOVA with Tukey’s multiple comparisons). **(D)** Quantification showed similar numbers of FISH-labeled bacteria adherent to sham-treated and bupivacaine-treated corneas after *S. aureus* inoculation. Data were normalized to baseline bacteria and expressed as the mean ± SD of bacteria per field of view. ns = Not Significant (Two-way ANOVA with Tukey’s multiple comparisons).

Next, enucleated WT eyes were inoculated with *S. aureus*. Quantification of FISH labeling following *S. aureus* inoculation and a 6 hour incubation *ex vivo* showed bacterial adhesion to wild-type corneas increased to match TRPV1 (-/-) corneas, after normalization to baseline as above. Thus, TRPV1-dependent corneal defense was abolished *ex vivo* (Fig. 2C).

Corneal sensory nerve firing was then blocked *in vivo* using sub-conjunctival bupivacaine which inhibits neuronal sodium channels ^42^. Bupivacaine treatment did not increase *S. aureus* adhesion to healthy WT mouse corneas *in vivo* versus sham-treated controls (Fig. 2D), i.e. corneas retained their inhibitory activity against *S. aureus* adhesion showing that nerve firing was not required. That result contrasted with our previous findings for *P. aeruginosa* in which bupivacaine treatment significantly increased bacterial adhesion to the mouse cornea ^25^. This latter result was reconfirmed in the present study (Supplemental Fig. S2).

To confirm TRPV1-specificity in corneal defense against *S. aureus* adhesion, we next used a selective TRPV1 antagonist JNJ-17203212 in WT mice (Fig. 3). This drug blocks neuronal TRPV1 activity both *in vivo* and *in vitro* by competing for the capsaicin binding site, rendering the channel inactive to other noxious stimuli ^38^. Local injection and topical application of JNJ-17203212 to WT corneas before *S. aureus* inoculation showed antagonistic effects lasting up to 3 hours in the murine cornea, demonstrated by decreased defensive wipes in a capsaicin eye wipe test (Fig. 3A). JNJ-17203212 treated corneas showed minimal fluorescein staining similar to controls indicating an intact epithelium (Fig. 3B) but showed a significant increase in *S. aureus* adhesion (> 20-fold) relative to controls (after normalization to baseline) (Fig. 3C), thus supporting a TRPV1-specific role in corneal defense against *S. aureus* adhesion.

**Figure 3.**
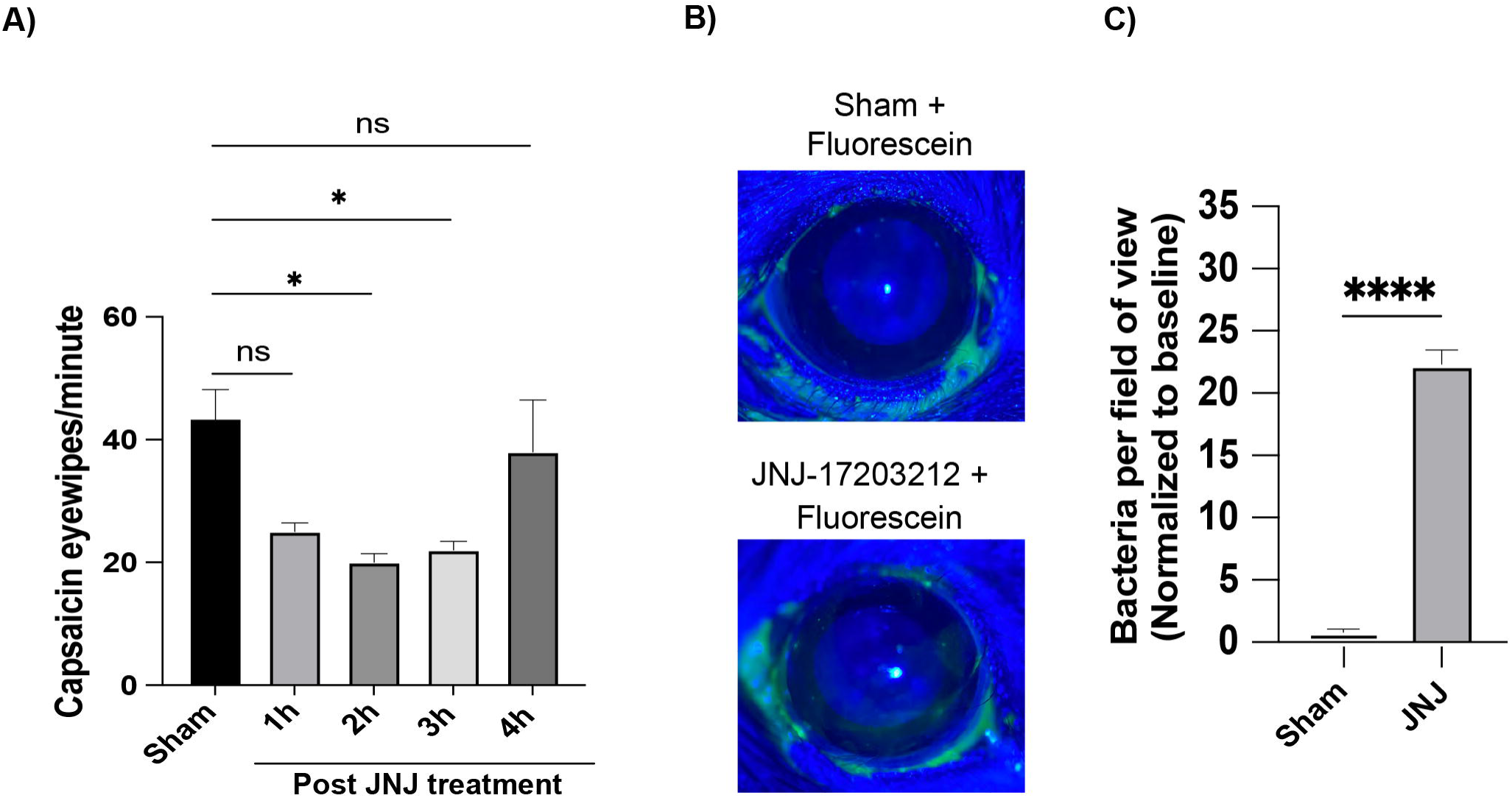
A TRPV1 antagonist JNJ-17203212 inhibits corneal defense versus *S. aureus.* **(A)** WT corneas treated with a TRPV1 antagonist JNJ-17203212 (500 µM) showed reduced sensitivity to capsaicin (100 µM) for 3 hours compared to sham controls. * P ˂ 0.05, ns = Not Significant (One-way ANOVA with Tukey’s multiple comparisons). **(B)** Slit-lamp images of WT corneas after TRPV1 antagonist treatment showed no corneal staining indicating an intact corneal epithelium. **(C)** Quantification showed a significant increase in FISH-labeled bacteria to antagonist treated WT corneas after *S. aureus* inoculation versus sham controls. Data were normalized to baseline bacteria and expressed as the mean ± SD of bacteria per field of view. **** P ˂ 0.0001 (Two-way ANOVA with Tukey’s multiple comparisons).

### *S. aureus* Challenge Does Not Elicit a Corneal CD11c^+^ Immune Cell Response

Previously we showed that increased *P. aeruginosa* adhesion to healthy or superficially-injured mouse corneas after RTX or bupivacaine treatment correlated with inhibition of corneal CD11c^+^ cell responses to *P. aeruginosa* ^25^. Prior to that study, we had also shown that corneal CD11c^+^ cell responses to *P. aeruginosa* correlated with the inhibition of bacterial adhesion to the cornea after superficial injury with immune cell movement in close proximity to attached bacteria ^21^. Thus, healthy cornea CD11c^+^ cell responses to *S. aureus* challenge were assessed using CD11c-YFP mice and compared to *P. aeruginosa* (Fig. 4). Results showed that *S. aureus* challenge did not cause a significant corneal CD11c^+^ cell response (Fig. 4A, B) contrasting with *P. aeruginosa* (Fig. 4F, G). CD11c^+^ cells also moved further away from the cornea surface (Fig. 4C) also in contrast with *P. aeruginosa* where migration towards the cornea surface was observed (Fig. 4H) as we had previously observed ^21^. Morphological analysis revealed CD11c^+^ cells became less spherical after *S. aureus* inoculation (Fig. 4D) with a similar result observed for *P. aeruginosa* (Fig. 4I), the latter consistent with our prior study ^25^. CD11c^+^ cell volume showed no after *S. aureus* inoculation (Fig. 4E) contrasting with increased cell volume observed after *P. aeruginosa* challenge (Fig. 4J) that was also consistent with our prior study ^25^. Thus, corneal CD11c^+^ cell responses to *S. aureus* were mostly distinct from those in response to *P. aeruginosa*, i.e. no change in cell number, cells moving in the opposite direction away from the corneal surface with no change in cell volume, suggesting a lack of CD11c^+^ cell involvement in defense versus *S. aureus* adhesion.

**Figure 4.**
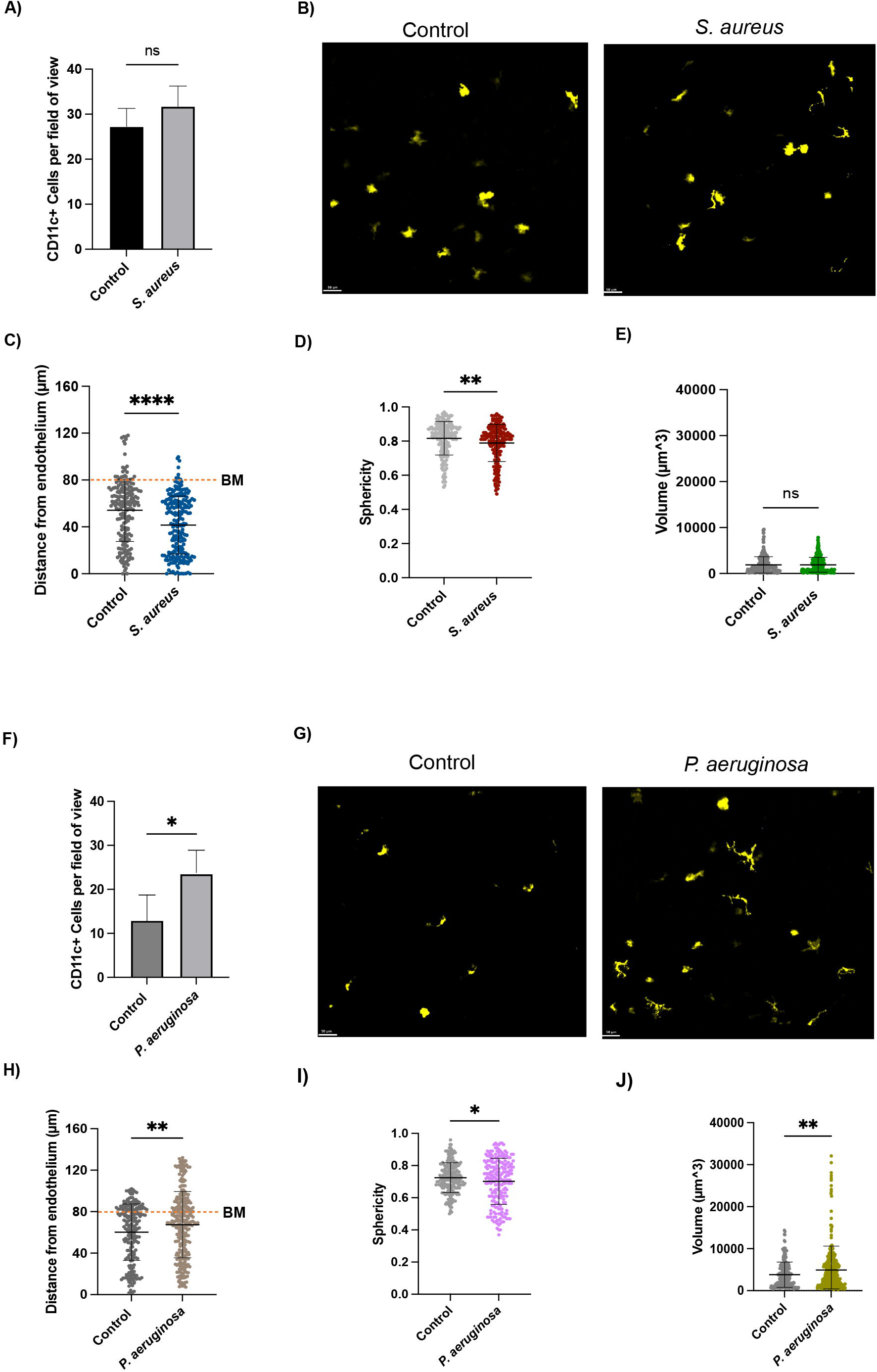
Distinct corneal CD11c^+^ cell responses to *S. aureus* compared to *P. aeruginosa.* **(A)** Quantification of CD11c^+^ cells in healthy WT corneas 4 hours after *S. aureus* challenge reveals no significant increase in cell numbers versus controls after inoculation. **(B)** Z-projections of the YFP channel showing all CD11c^+^ cells (yellow) projected into one plane. Scale bars = 50 µm. **(C)** CD11c^+^ cells were further away from the corneal epithelium after *S. aureus* inoculation. The dashed line denotes the basement membrane (BM) with areas above representing the epithelium and zero indicating the endothelium. **(D)** CD11c^+^ cells were significantly less spherical after *S. aureus* inoculation. **(E)** CD11c+ cells showed no change in volume after *S. aureus* inoculation. **(F)** Quantification of CD11c^+^ cells in healthy WT corneas 4 hours after *P. aeruginosa* challenge shows a significant increase in CD11c^+^ cells versus controls. **(G)** Z-projections of the YFP channel showing all CD11c^+^ cells (yellow) projected into one plane. Scale bars = 50 µm. **(H)** CD11c^+^ cells were significantly closer to the corneal epithelium following *P. aeruginosa* inoculation. The dashed line denotes the basement membrane (BM) as above. **(I)** CD11c^+^ cells were less spherical after *P. aeruginosa* inoculation. **(J)** CD11c^+^ cells showed an increased volume after *P. aeruginosa* inoculation. Data were expressed as the mean ± SD cells per field of view. * P ˂0.05, ** P ˂ 0.01, **** P ˂ 0.0001, ns = Not Significant (Student’s t-Test).

### *S. aureus* Challenge Did Not Elicit a Corneal Lyz2^+^ Immune Cell Response

Given the absence of CD11c^+^ cell responses to *S. aureus*, we expanded our investigation to include a different subset of corneal immune cells expressing Lyz2, e.g. monocytes, macrophages. Inoculation of mT/mG + LysMcre hybrid WT mice with *S. aureus* did not result in an increase in Lyz2^+^ cells (Fig. 5A, B). Interestingly, the same result was found for *P. aeruginosa* (Fig. 5F. G) contrasting with the observed CD11c^+^ cell response (Fig. 4F, G). There was no change in Lyz2^+^ cell migration in response to *S. aureus* (Fig. 5C) contrasting with the migration of CD11c^+^ cells away from the surface under the same conditions (Fig. 4C). However, *P. aeruginosa* challenge did result in Lyz2^+^ cell migration towards the corneal epithelium (Fig. 5H) as observed for CD11c^+^ cells (Fig. 4H). While no change in cell numbers was observed, Lyz2^+^ cell morphologies changed after *S. aureus* inoculation with increased Lyz2^+^ cell sphericity (Fig. 5D) and volume (Fig. 5E) contrasting with CD11c^+^ cell changes (Fig. 4D, E). *P. aeruginosa* challenge did not impact Lyz2^+^ cell sphericity (Fig. 5I) or volume (Fig. 5J) contrasting with CD11c^+^ cell changes (Fig. 4I, J).

**Figure 5.**
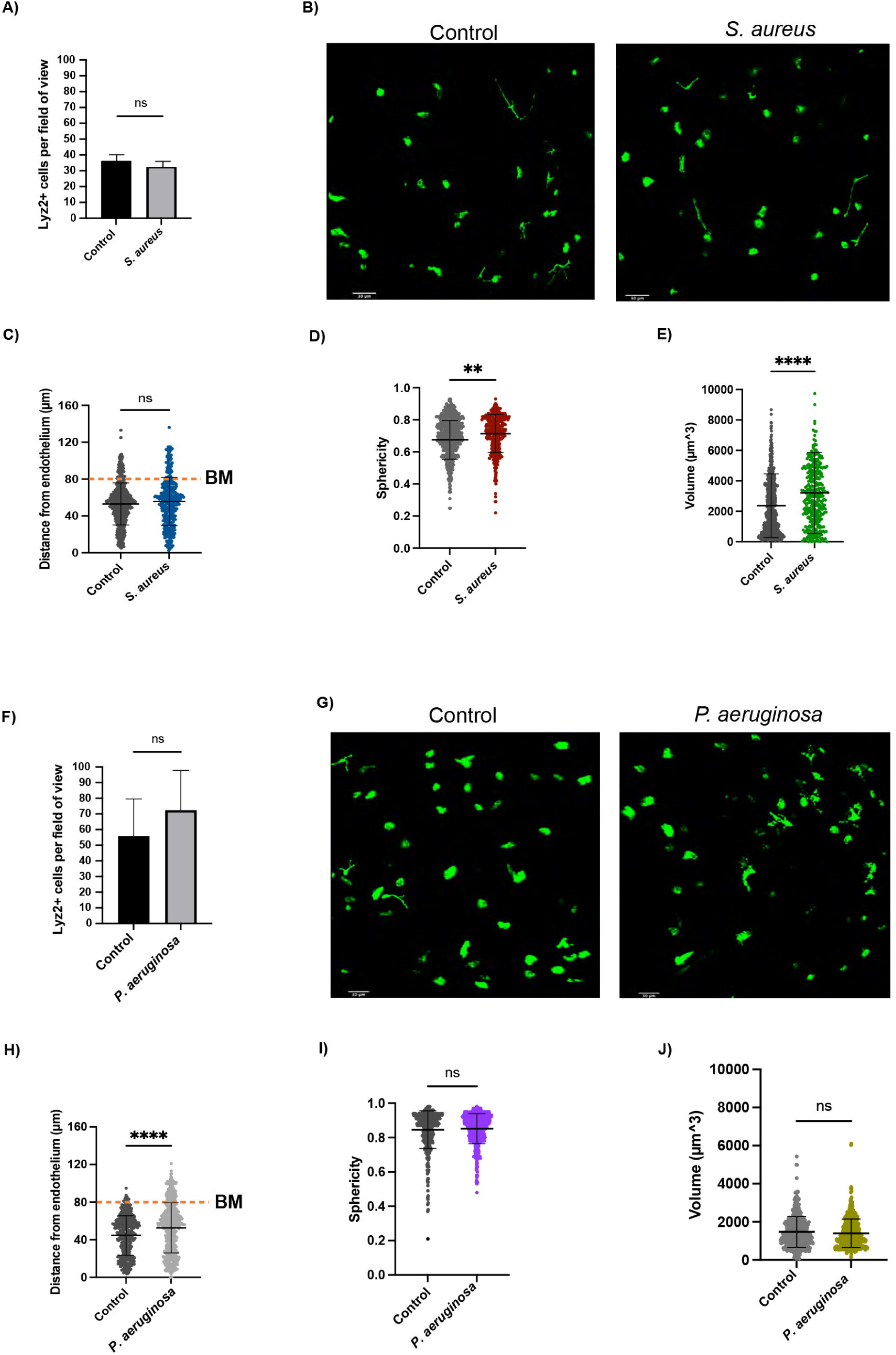
Quantitative and morphological analysis of Lyz2^+^ cells in healthy corneas of mT/mG + LysMcre (red cell membranes; green Lyz2^+^ immune cells) hybrid WT mice at 4 hours after *S. aureus* or *P. aeruginosa* challenge. **(A)** Quantification of Lyz2^+^ cells after *S. aureus* challenge showing no change in cell numbers. **(B)** Z-projections of the GFP channel with all Lyz2^+^ cells (green) projected into one plane. Scale bars = 30 µm. **(C)** Lyz2^+^ cells did not show migration changes after *S. aureus* inoculation. The dashed line denotes the basement membrane (BM) with areas above representing the epithelium and zero indicating the endothelium. **(D)** Lyz2^+^ cells were more spherical after *S. aureus* inoculation, and **(E)** showed a significant increase in cell volume. **(F)** Quantification of Lyz2^+^ cells after *P. aeruginosa* challenge shows no significant increase in cell numbers. **(G)** Z-projections of the GFP channel with all Lyz2^+^ cells projected into one plane. Scale bars = 30 µm. **(H)** Lyz2^+^ cells were significantly closer to the corneal epithelium following *P. aeruginosa* inoculation. The dashed line denotes the basement membrane (BM) as above. **(I)** Lyz2^+^ cell sphericity did not change after *P. aeruginosa* inoculation. **(J)** Lyz2^+^ cell volume did not change after *P. aeruginosa* inoculation. Data expressed as the mean ± SD cells per field of view. ** P ˂ 0.01,**** P ˂ 0.0001, ns = Not Significant (Student’s t-Test).

### Corneal CD45+ Cells Respond to *S. aureus* Challenge

The absence of corneal CD11c^+^ or Lyz2^+^ cell infiltrative responses to *S. aureus* led us to expand our investigation to include CD45^+^ cells using antibody labeling in healthy WT mice. Results showed a small but significant increase in corneal CD45^+^ immune cells after *S. aureus* inoculation (∼1.7-fold) which was inhibited by RTX (Fig. 6A), showing that this response required TRPV1-expressing sensory nerves, and also consistent with our previous work with *P. aeruginosa* ^25^. Morphology analysis showed that CD45^+^ cells became more spherical and following *S. aureus* inoculation (Fig. 6B) as observed for Lyz2^+^ cells (Fig. 5D). That change was reversed by RTX treatment with cells becoming more dendriform (Fig. 6B).

**Figure 6.**
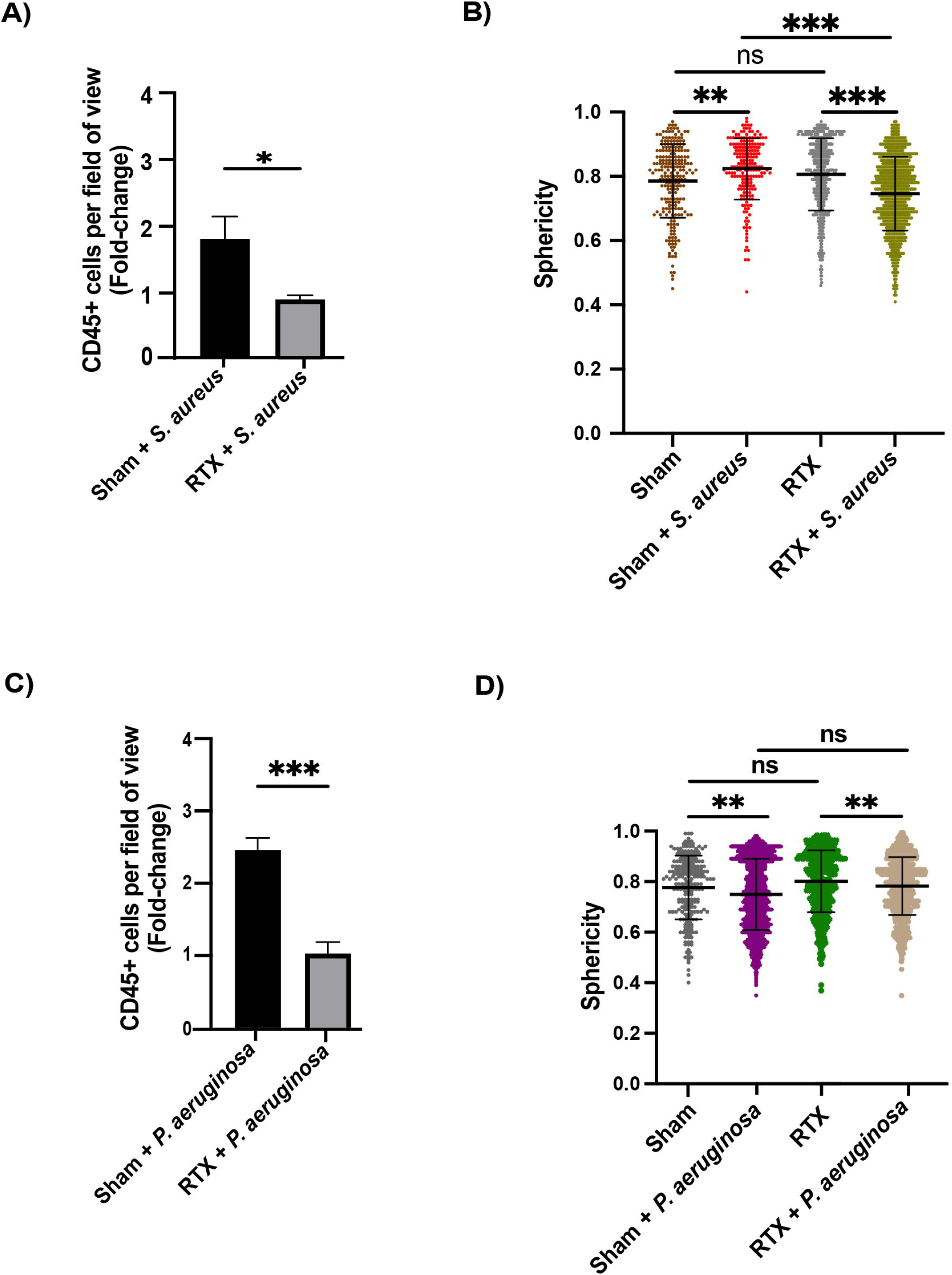
Increased CD45^+^ cell numbers at 4 hours after *S. aureus* inoculation was blocked by RTX. **(A)** Quantification of CD45^+^ cells reveals a small but significant increase in CD45+ cells after *S. aureus* inoculation (∼1.7-fold) which was inhibited by RTX. Data normalized to baseline CD45+ cells for each group. **(B)** Morphology analysis of corneal CD45+ cells showed cells were more spherical after *S. aureus* inoculation, a change reversed by RTX. **(C)** Quantification of corneal CD45+ cells reveals a significant increase after *P. aeruginosa* inoculation (∼2.5-fold) that was blocked by RTX. Data normalized to baseline CD45+ cells for each group. **(D)** Morphological analysis of CD45^+^ cells shows a reduced sphericity after *P. aeruginosa* inoculation that was not affected by RTX. Data were expressed as a mean ± SD cells per field of view. * P ˂ 0.05, ** P ˂ 0.01, *** P ˂ 0.001, **** P ˂ 0.0001, ns = Not Significant (Student’s t-Test).

We had previously shown that the healthy cornea CD45+ response to *P. aeruginosa* challenge was abrogated in TRPA1/TRPV1 (-/-) mice ^25^. Here, we confirmed sensory nerve dependence of that response using RTX (Fig. 6C). CD45^+^ cell morphological changes in response to *P. aeruginosa* showed that cells became less spherical, a change unaffected by RTX (Fig. 6D). This change in sphericity contrasted with the CD45^+^ cell response to *S. aureus* (Fig. 6B), which did respond to RTX. Therefore, while both *S. aureus* and *P. aeruginosa* induced a nerve-dependent CD45^+^ cell response, morphological responses of these cells differed between these pathogens.

## Discussion

We have previously shown that the unique ability of the healthy murine cornea to resist adhesion by *P. aeruginosa*^2^, and to resist bacterial microbiome formation^1^, involves corneal sensory nerves expressing Transient Receptor Potential (TRP) ion channels: TRPA1 required for defense against *P. aeruginosa* and TRPV1 to defend against the adhesion of environmental bacteria^25^. Here, we tested if there was pathogen specificity in this TRP-mediated corneal defense versus bacterial adhesion by using *S. aureus*, a different ocular pathogen. Interestingly, results showed that corneal resistance to *S. aureus* adhesion required corneal sensory nerves expressing TRPV1. This conclusion was derived from a combination of experiments firstly using TRPV1 (-/-) mice, followed by RTX ablation of TRPV1-expressing sensory nerves in WT mice, then testing intact WT corneas *ex vivo*. Since RTX also ablates the subset of TRPV1-expressing nerves that co-express TRPA1 ^41^, specificity for TRPV1 in defense versus *S. aureus* was confirmed using the selective antagonist JNJ-17203212. In contrast to our previous findings with *P. aeruginosa*, bupivacaine treatment of healthy WT corneas did not compromise defense versus *S. aureus* adhesion, showing that sensory nerve firing was not required. Subsequent studies of immune cell responses to *S. aureus* challenge, with comparison to *P. aeruginosa*, showed differences in immune cell types that responded to each pathogen: Lyz2^+^ cells for *S. aureus*, versus CD11c^+^ cells for *P. aeruginosa* ^25^ (also confirmed in this study). While both pathogens induced sensory nerve-dependent CD45^+^ cell responses, their morphology changes differed between the two pathogens.

Like *P. aeruginosa, S. aureus* poses a significant threat to ocular health, particularly in the context of corneal infections, which can lead to vision impairment ^5,43,44^. It is interesting, therefore, that TRPV1 (and not TRPA1) was required for defense against *S. aureus*, in direct contrast to the role of TRPA1 (and not TRPV1) in countering *P. aeruginosa* ^25^. The basis for this selectivity is not yet known. It is possible that TRPV1-expressing sensory nerves detect cell wall antigens of Gram-positive bacteria such as *S. aureus*, and a subset of TRPA1-expressing nerves (also TRPV1 positive ^41^) detect Gram-negative bacterial cell wall antigens, each responding accordingly to limit adhesion. In that regard, present findings are consistent with our previous work showing a role for TRPV1 in defending the cornea against adhesion of environmental bacteria, which are thought to be mostly Gram-positive, and which could include conjunctival commensals e.g. coagulase-negative *Staphylococcus* and *Propionibacterium* spp. ^45^. Furthermore, other prior studies have shown that *P. aeruginosa* LPS promotes inflammation and pain *via* TRPA1 activation ^46^, and *S. aureus* pore-forming toxins such as α-hemolysin can trigger inflammation and pain *via* TRPV1-expressing neurons ^31^. However, TRPA1 and TRPV1 can also both bind bacterial-derived quorum-sensing molecules to activate these channels (Gram-negative-derived molecules) or show inhibitory effects (Gram-positive-derived molecules) with differences in TRP receptor specificity depending upon quorum-sensing molecular structure ^47^. It is also possible that selectivity between TRPV1-and TRPA1-mediated defenses is derived from differences in bacterial susceptibility to effector mechanisms triggered by various bacterial antigens, e.g. if *P. aeruginosa* resists TRPV1-triggered defenses while *S. aureus* is susceptible. The molecular basis for TRPV1 vs. TRPA1 selectivity in corneal defense against bacterial adhesion is a focus of our ongoing studies. Our findings to date suggest, however, that TRP selectivity is not based on bacterial categorization as a pathogen or an environmental/commensal but could be based upon differences in cell wall composition.

TRPV1 is expressed on the majority of corneal sensory neurons ^48^. While some studies indicate TRPV1 expression on non-neuronal cell types ^49–51^, the present study suggests that TRPV1 defense versus *S. aureus* adhesion involves nerve-associated TRPV1. This conclusion reflects an inhibition of this corneal defense by RTX, which ablates TRPV1-expressing sensory nerves, and the loss of this defense when eyes were tested *ex vivo*. Contrasting with *P. aeruginosa* ^25^, however, corneal defense versus *S. aureus* adhesion was retained after bupivacaine block of sensory nerve firing. These findings suggesting a fundamental difference between mechanisms of nerve-associated TRPV1-and TRPA1-mediated corneal defenses versus *S. aureus* and *P. aeruginosa* respectively.

Effector mechanism(s) by which TRPV1 defends the cornea versus *S. aureus* adhesion remain to be determined. The lack of requirement for sensory nerve firing suggests TRPV1 allows a local release of factors with antimicrobial or other anti-adhesive efficacy. Effectiveness of a TRPV1 antagonist JNJ-17203212 in blocking corneal defense versus *S. aureus* adhesion suggests that TRPV1 channel activation with Ca^2+^ influx is required ^49,52^ since these would be blocked by this competitive antagonist. Moreover, activation of TRPV1 by agonists, bacterial ligands or other potentially harmful stimuli can result in the local release of neuropeptides, e.g. Calcitonin-Gene-Related Peptide or Substance P, with subsequent pro-inflammatory ^53^, antimicrobial ^54^, or immunomodulatory effects ^55^ which could all contribute to corneal defense ^56^. Importantly, while activation of TRPV1 with Ca^2+^ influx (i.e. membrane depolarization) is associated with generation of an action potential and sensory nerve firing ^57^, neuropeptide(s) release independently of those events could provide an avenue to explain the retention of TRPV1-mediated defense against *S. aureus* adhesion despite bupivacaine block of sensory nerve firing *via* inhibition of voltage-gated sodium channels ^58^. Indeed, TRPV1 can control neuronal release of the neurotransmitter glutamate independently of action potential evoked release, albeit with both under Ca^2+^ regulation ^59^.

It is well-established that TRPV1 and TRPA1 activation has multiple effects on immune cells in many tissues ^28,55,60^. Previously we showed that TRPA1-mediated defense against *P. aeruginosa* adhesion to healthy and superficially-injured corneas closely correlated with changes in immune cell recruitment (CD45+ and CD11c^+^ cells) and morphology (CD11c^+^ cells) ^25^. We also showed that TRPV1 and TRPA1 contributed to baseline levels of resident corneal immune cells, e.g. MHC-II positive cells, and to contact lens-induced corneal parainflammatory responses ^37^. The present study showed distinct differences in immune cell responses to *S. aureus* compared to *P. aeruginosa*. The most obvious was absence of a quantitative CD11c^+^ cell response to *S. aureus*, i.e. no change in cell number. CD11c^+^ cells also migrated away from the corneal epithelium after *S. aureus* challenge and showed no change in cell volume. Other than a reduction in cell sphericity after *S. aureus* challenge, all contrasted with CD11c^+^ responses to *P. aeruginosa* suggesting that corneal CD11c^+^ cells, e.g. resident dendritic cells, may play a less significant role in defending the cornea against *S. aureus* adhesion compared to *P. aeruginosa*.

Lyz2^+^ cells, however, responded to *S. aureus*, not quantitatively but with morphology changes of increased sphericity and volume in response to the bacteria compared to the absence of Lyz2^+^ cell changes, quantitative or morphological, in response to *P. aeruginosa*. Thus, corneal responses to *S. aureus* challenge appear distinct from those in response to *P. aeruginosa*: the former favoring Lyz2^+^ cells, the latter favoring CD11c^+^ cells not expressing Lyz2. Ongoing flow cytometric analysis of immune cell responses to these two bacterial pathogens will help define Lyz2^+^ and CD11c^+^ cell types involved. Complementary studies will then determine the functional role(s) of identified cells in defending the cornea against bacterial adhesion.

*S. aureus* challenge was found to induce a small but significant quantitative corneal CD45^+^ cell response that was inhibited by RTX suggesting the requirement of TRPV1-expressing sensory nerves. This was consistent with *P. aeruginosa* induced quantitative corneal CD45^+^ responses that we reported previously using TRPA1/TRPV1 double gene knockout mice ^25^ and confirmed here using RTX. Thus, corneal CD45^+^ cells respond to both *S. aureus* and *P. aeruginosa* challenge in a sensory nerve-dependent manner. Further studies are needed to determine the specific type(s) of CD45^+^ cells responding to each pathogen and the proportion of these cells also expressing Lyz2 or CD11c. Differences in CD45^+^, Lyz2^+^ and CD11c^+^ cell morphology changes in response to both pathogens may reflect differences in cellular activation states and respective involvement of TRPV1-versus TRPA1-expressing sensory nerves. However, further studies will be needed to determine the significance of these cellular responses (quantitative, directional and morphological) with respect any role(s) in defending the healthy cornea against bacterial adhesion.

In conclusion, this study shows that TRPV1-expressing sensory nerves are required for defending the healthy cornea against the adhesion of *S. aureus*, an important ocular and systemic pathogen. This defense is independent of sensory nerve firing. These findings contrast with *P. aeruginosa* for which TRPA1 and sensory nerve firing are required. While specific mechanisms of TRPV1-mediated defense versus *S. aureus* remain to be determined, a role for CD45^+^ and Lyz2^+^ cells cannot be excluded. Corneal immune cell responses to *S. aureus* shown here appear distinct from the sensory nerve-dependent CD11c^+^ cell responses that correlate with TRPA1-mediated defense against *P. aeruginosa* ^25^. Ongoing studies are focused on determining the antimicrobial or anti-adhesive mechanisms underlying TRPV1-and TRPA1-mediated sensory nerve defense against these two clinically significant ocular pathogens, and the role of specific immune cell types associated with these responses.

## Acknowledgements

This work was supported by the National Institutes of Health EY030350 (SMJF).

## Supplemental Material

**Supplemental Figure S1.** Morphological analysis of corneal immune cells using Imaris. **(A) Left image**: A “surface object” of each immune cell (representative image shows Lyz2^+^ immune cells, green) was created using a XY size of 10 µm. **Center image:** Another surface object was manually drawn using the red channel (or reflectance for CD11c^+^-YFP mice) to select the total immune cells signal above the corneal endothelium (pink). **Right image**: From the created endothelium, a distance transformation was performed. This created a new channel (channel 3, not shown in the image) to determine the distance of Surface 1 (Immune cells) from Surface 2 (Endothelium). Scale bars = 30 µm. **(B) Left image**: Immune cells were statistically color-coded to reflect sphericity ranges: Purple ≤ 0.3 = most dendriform, 0.3˂ ˃0.8, and red ≥ 8 = most circular. **Center image**: Immune cells were statistically color-coded to reflect ranges in volumes/sizes. **Right image:** Immune cells were assessed for their location in the cornea based on their mean intensity (distance) from the endothelium. Cells were statistically coded to reflect ranges in distance: Purple, 0 µm = at the endothelium, 0˂ ˃80 µm = stroma, and red ≥ 80 µm = epithelium. Scale bars = 30 µm.

**Supplemental Figure S2.** Increased *P. aeruginosa* adherence to wild-type heathy mouse corneas after bupivacaine treatment. **(A)** Corneas were labeled with a universal 16S rRNA-targeted FISH probe to detect background environmental bacteria in uninoculated controls and *P. aeruginosa* after inoculation. Representative images show *P. aeruginosa* inoculated corneas with increased bacterial adhesion to corneas treated with bupivacaine (0.5%) versus sham controls following hourly inoculation of ∼10^11^ CFU/ml *P. aeruginosa* for 4 hours. Scale bars = 50 μm. Only bacterial channel shown. **(B)** Quantification of FISH-labeled bacteria on wild-type corneas after *P. aeruginosa* inoculation showing a ∼ 2.6-fold increase in bacterial adhesion to bupivacaine treated corneas versus sham controls. Data were normalized to environmental (baseline) bacteria and expressed as a mean ± SD of bacteria per field of view. * P < 0.05 (One-way ANOVA with Tukey’s multiple comparisons).

**Supplemental Video 1**. Video shows how the surfaces of the corneal endothelium and immune cells were rendered using the Imaris surface function. A 3D volume image of the central cornea from a mT/mG + LysMcre hybrid WT mouse (red cell membranes; green Lyz2^+^ immune cells) is shown. A white surface object was generated from the endothelium signal, and the distance outside this object was transformed into a new channel, where intensity values (from low to high) indicate increasing distance from the endothelium. Then, the green surface objects were generated from the Lyz2^+^ signal, and distance of these cell objects from the endothelium object was measured according to the intensity values of the transformed channel.

**Supplemental Video 2**. Video shows how surface rendered immune cells can be categorized by different properties. Firstly, the immune cell objects were color-coded by their volume, with a colormap in the lower right corner ranging from purple (smallest) to red (largest). Next, the cell objects were differently color-coded by their sphericity, from purple being most dendriform shape to red being the most spherical. Finally, the cell objects were color-coded by their distance from the endothelium object, from purple being closest to red being farthest. The image was also rotated to better visualize spatial distribution of the cells.

