## Supplementary figures and images for "TRPV1 Defends the Healthy Murine Cornea against *Staphylococcus aureus* Adhesion Independently of Sensory Nerve Firing"

### Supplemental Figure S1

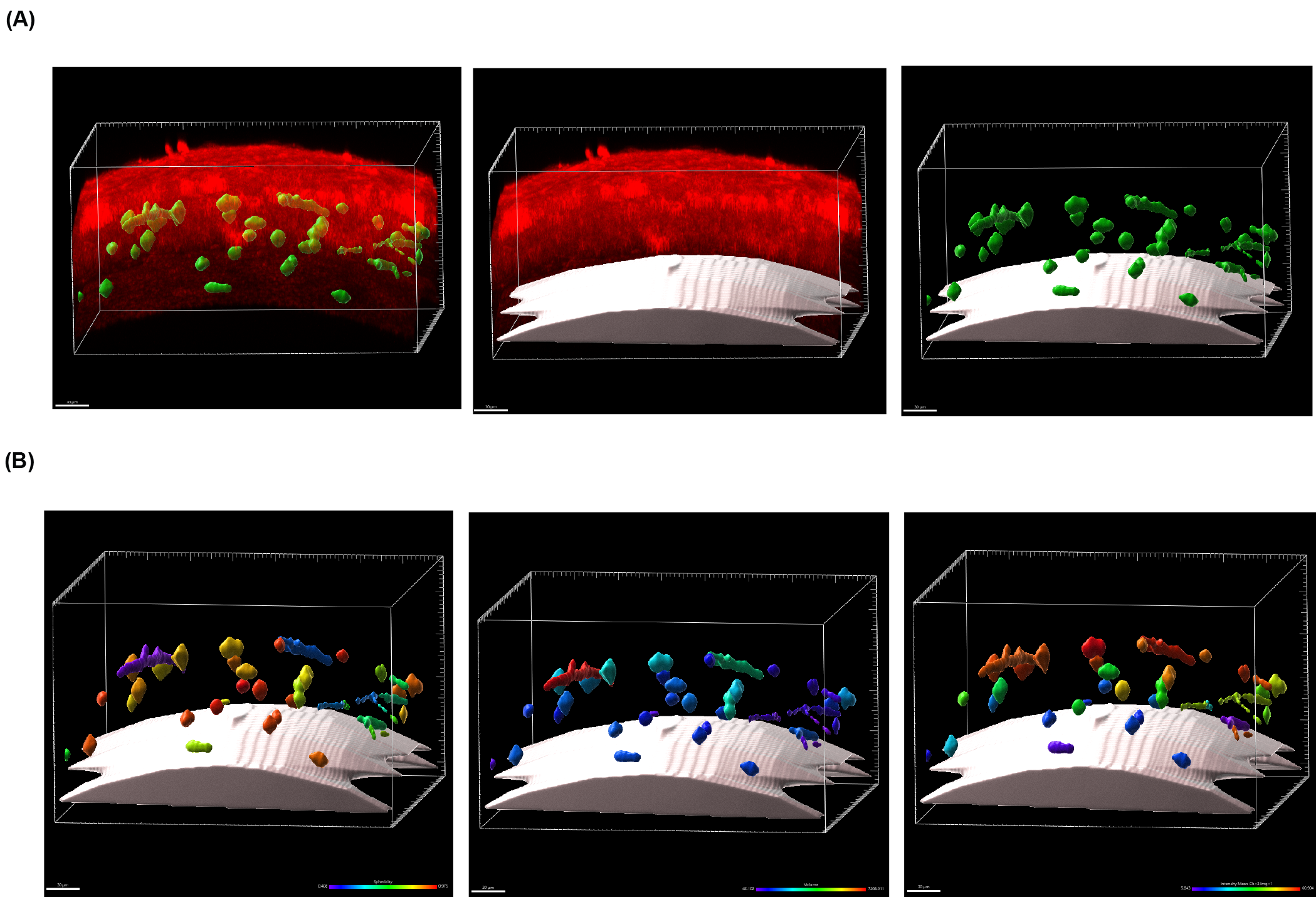

### Supplemental Figure S2

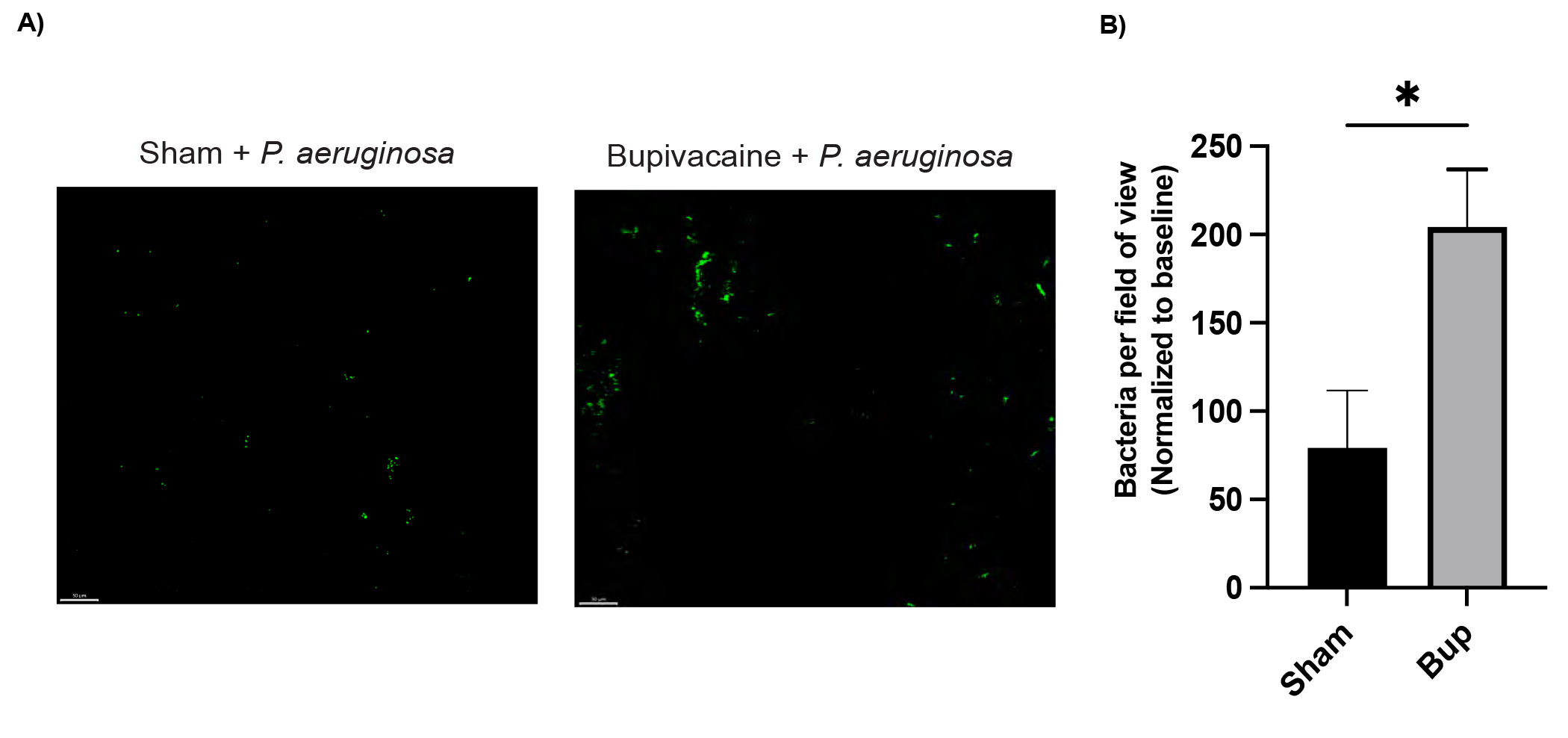
